# Overcoming systematic errors caused by log-transformation of normalized single-cell RNA sequencing data

**DOI:** 10.1101/404962

**Authors:** Aaron Lun

**Affiliations:** Cancer Research UK Cambridge Institute, University of Cambridge, Li Ka Shing Centre, Robinson Way, Cambridge CB2 0RE, United Kingdom

## Abstract

Applying a log-transformation to normalized expression values is one of the most common procedures in exploratory analyses of single-cell RNA sequencing (scRNA-seq) data. Normalization removes systematic biases in sequencing coverage between cells, while the log-transformation ensures that downstream computational procedures operate on relative rather than absolute differences in expression. We show that the log-transformation can introduce systematic errors when cells vary in sequencing coverage, leading to spurious non-zero differences in expression and artificial population structure in simulations. We observe similar effects in real scRNA-seq data where the difference in transformed values between groups of cells is not an accurate proxy for the log-fold change. We provide some practical recommendations to overcome this effect and analytically derive an expression for a larger pseudo-count that controls the transformation-induced error to a specified threshold.

## 1 Background

Log-transformed expression values are widely used in exploratory analyses of single-cell RNA sequencing (scRNA-seq) data. This is driven by the simplicity of the log-transformation and the ease of interpretation of the transformed values. In particular, differences between log-values can be used as proxies for log-fold changes in expression between cells, which are often more relevant than differences in the counts. This is useful for ensuring that relative rather than absolute differences in the counts are used in distance-based procedures like clustering or trajectory construction. The log-transformation also reduces the impact of stochastic fluctuations in the counts for high-abundance genes. This would otherwise result in large differences between cells that are mostly uninteresting as the fold changes between counts are small.

Despite its popularity, the log-transformation has a number of problems such as suboptimal variance stabilization and arbitrariness in the choice of pseudo-count. One issue of particular interest is that the mean of the log-counts is not generally the same as the log-mean count [1]. This is problematic in scRNA-seq contexts where the log-transformation is applied to normalized expression data. Here, normalization is performed to remove inter-sample biases on the count scale [2, 3]. For genes that are not differentially expressed (DE), the expectation of the normalized expression is the same between cells. However, the expectation of the log-transformed normalized values may not be the same, resulting in spurious differences between cells on the log-scale.

The ideal solution would be to use methods that model the counts directly to account for the mean-variance relationship. For example, we could use *BASiCS* [4] for detecting highly variable genes or *zinbwave* [5] for factor analysis. This would avoid the need for any transformation and subsequent distortion of the expected expression. However, most existing methods for scRNA-seq data analysis (e.g., clustering, dimensionality reduction) do not use count-based models. This is due to the difficulties in estimating model parameters, often separately for each gene and/or each cell, compared to algorithms that only need distances between cells as input. Thus, it is still desirable to obtain transformed data for compatibility with existing methods.

Here, we describe the nature and impact of the discrepancy between log-mean and mean-log values in real and simulated data. We suggest some practical strategies to protect against this effect, including the use of DE analyses with count-based models as validation. We also derive an expression for a larger pseudo-count that can be added prior to transformation to control the magnitude of the discrepancy for each gene.

## 2 Quantifying the discrepancy

Let *X*_*ig*_ denote a random variable for a non-negative count of a gene *g* in cell *i*. The size factor *s*_*i*_ represents the scaling bias in *i*, which is removed by computing the normalized expression value *X*_*ig*_*/s*_*i*_ [6]. The log-transformed normalized expression is defined as

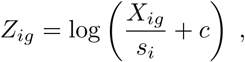

where *c* is a positive “pseudo-count” that ensures that the transformation is defined for *X*_*ig*_ *≥* 0 and *s*_*i*_ *>* 0. This calculation and its variants are widespread in analyses of count data from high-throughput sequencing experiments – for example, a similar approach is used by the cpm function in the *edgeR* package [7], differing only in a slight adjustment to the effective library size (equivalent to *s*_*i*_) for a given prior count (equivalent to *c*). The second-order Taylor series approximation for the expectation of *Z*_*ig*_ is

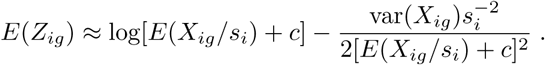

This expectation represents the mean of the log-normalized expression, while the first term on the right represents the log-mean normalized expression (after adding *c*). The second term on the right represents the discrepancy between these two values.

Now, consider two cells *i* = 1 and *i* = 2 that differ in their *s*_*i*_. Assume that *g* is non-DE between these two cells such that *E*(*X*_*ig*_*/s*_*i*_) = *µ*_*g*_ for both. The true log-fold change in the normalized expression values between these two cells is

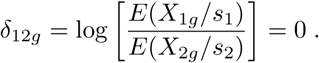

In analyses based on the log-transformed values, we use the expected difference in *Z*_*ig*_ as a proxy for *δ*_12*g*_. First, we simplify the expression for *E*(*Z*_*ig*_) to

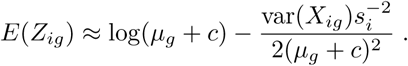

This means that the difference in *E*(*Z*_*ig*_) between these two cells is

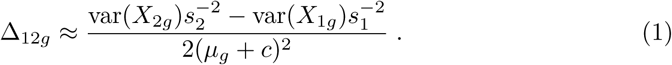

In general, Δ_12*g*_ ≠ 0 due to differences in the size factors and subsequently variances (driven by the strong mean-variance relationship in count data). This represents a spurious difference on the log-scale as the gene is not actually DE, i.e., *δ*_12*g*_ = 0.

We performed simulations to quantify this spurious difference in a range of scenarios for the typical *c* = 1 (Figure 1). We observed that it was most prominent when the size factors were different and either or both of them were less than unity. This is consistent with Equation 1, where Δ_12*g*_ increases in magnitude with smaller *s*_*i*_. We note that a 10-fold difference in the size factors across cells is not unusual in real scRNA-seq data [3] due to the variability of capture efficiency and amplification. In this situation, we can easily obtain a difference of 1 in the mean log_2_-values between cells. This corresponds to an artificial 2-fold change in expression that does not correspond to any biological effect.

**Figure 1.**
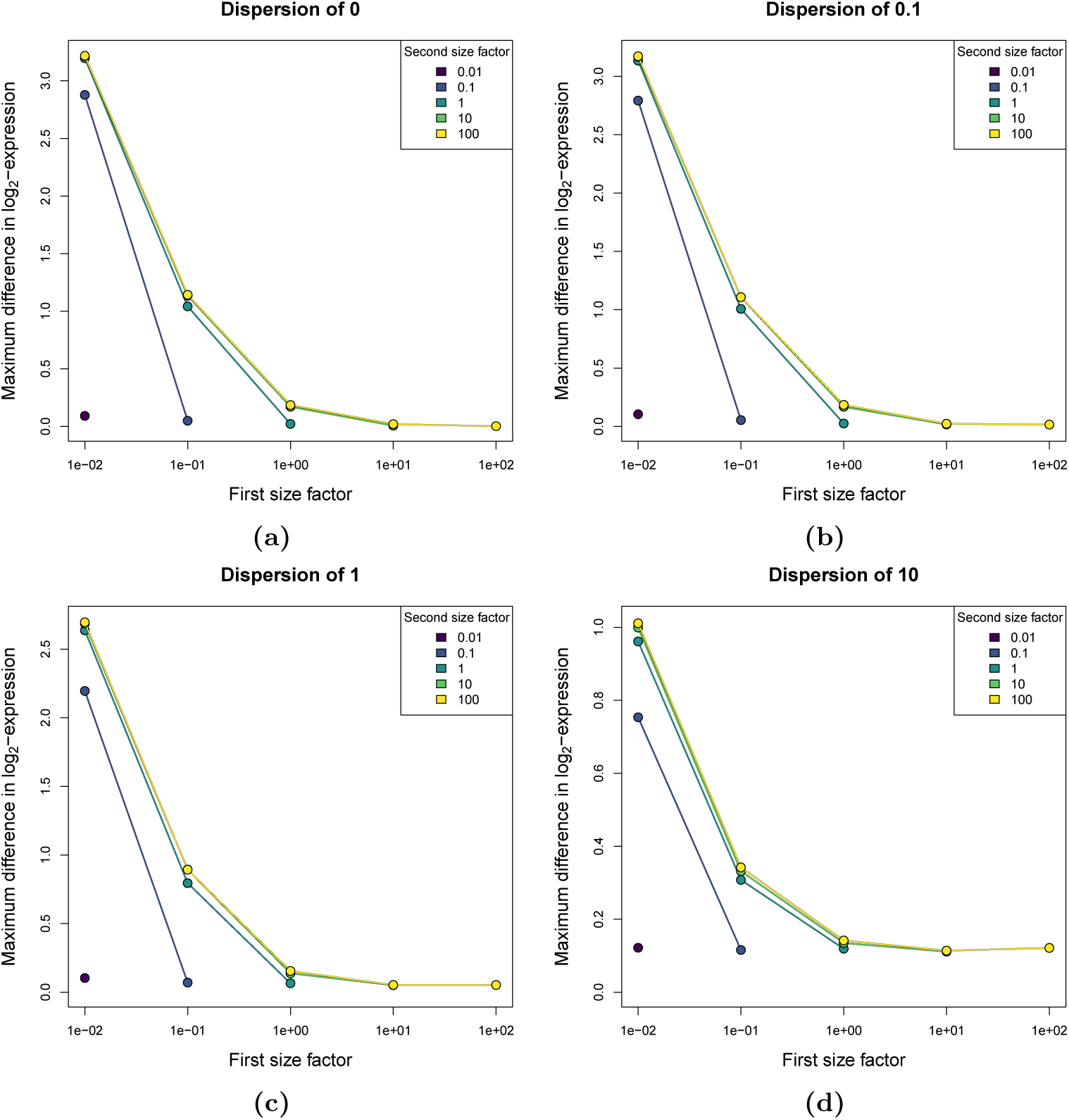
Maximum difference in the mean log_2_-expression values for a non-DE gene due to size factor differences. We simulated counts for non-DE genes in two groups of cells, where the size factor was the same for all cells in the same group (see Methods). Each simulation scenario was defined by the size factor used for the first group (*a*_1_), that for the second group (*a*_2_), and the negative binomial dispersion. For each scenario, we computed the difference between the mean *Z*_*ig*_ for each group and determined the maximum difference across genes of varying abundance. Values represent the average of the maximum across 10 simulation iterations. Standard errors were negligible and are not shown. For simplicity, we only show results for scenarios where *a*_1_ *≥ a*_2_.

We verified the existence of this effect in real scRNA-seq data from the 10X Genomics platform [8]. We used the publicly available ERCC data set (see Methods) in which gel emulsion beads containing ERCC spike-in transcripts were captured into droplets. The composition of the ERCC spike-in set should be constant in all droplets, so there should not be any difference in the expression profiles after library size normalization. However, we observed a systematic non-zero difference in the mean log-expression values when comparing the smallest and largest droplets (Figure 2a). This is fully attributable to the log-transformation, as no such difference is present in the log-fold change between the mean normalized expression values (Figure 2b). Discrepancies between the difference in mean log-values and the log-fold change are also present in real scRNA-seq data sets with variable size factors (Figures 2c, d).

**Figure 2.**
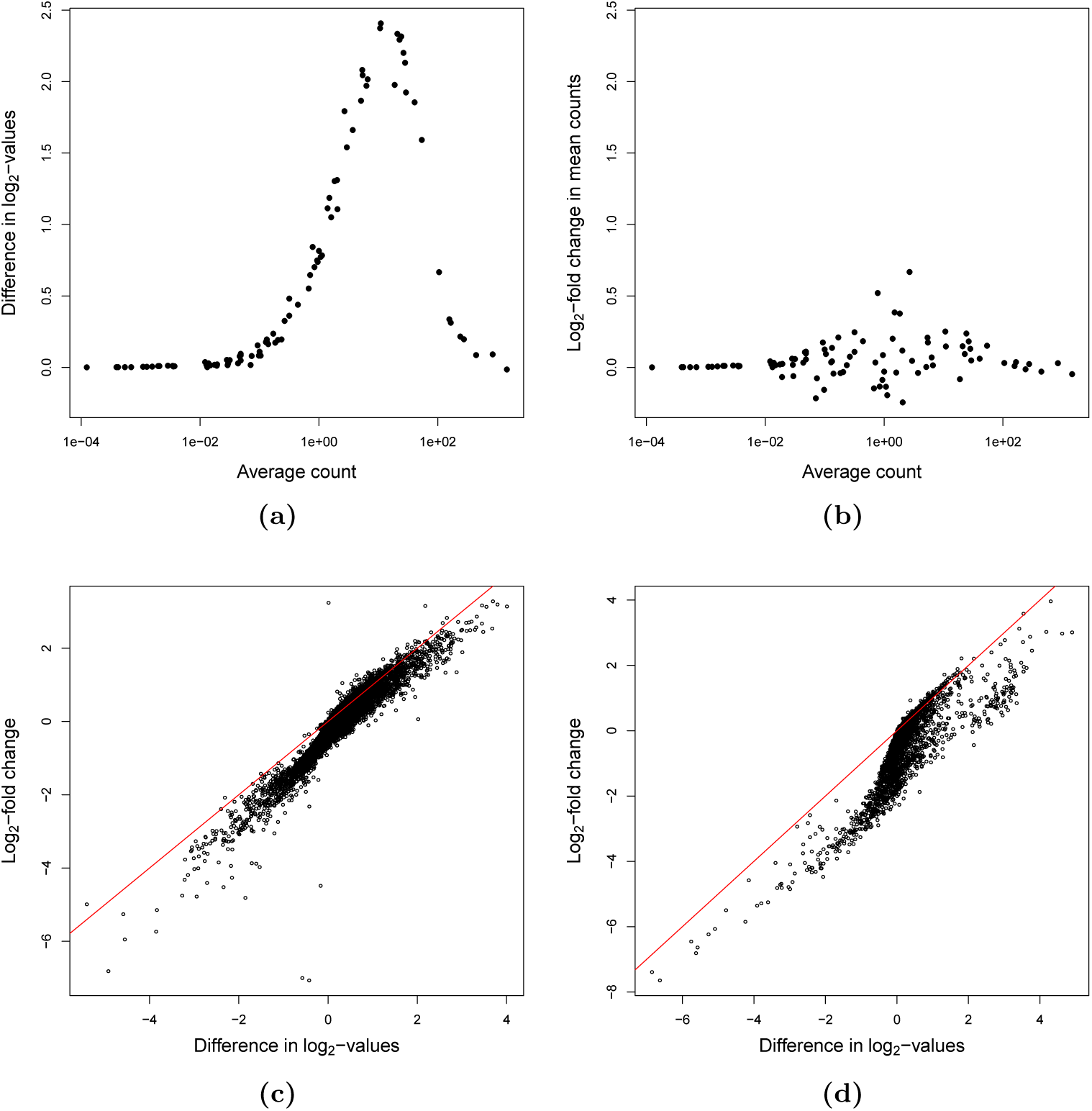
Effect of the log-transformation on comparisons between groups in real scRNA-seq data. (a) Differences in the mean *Z*_*ig*_ between the top 20% of libraries with the largest size factors and the bottom 20% in the ERCC 10X Genomics data set (see Methods). Each point represents a single ERCC spike-in transcript. (b) Log-fold change in the mean normalized expression between the same groups of libraries for each ERCC transcript. (c) Differences in the mean *Z*_*ig*_ for two clusters of cells with the largest and smallest median size factors in a brain scRNA-seq data set [9], compared to the log-fold change computed from the mean normalized expression. Each point represents a gene. Same as (c) for the peripheral blood mononuclear cell (PBMC) data set [8].

The effect of spurious differences in *E*(*Z*_*ig*_) is amplified in procedures that compute distances between cells. Consider the Euclidean distance between the expected log-normalized expression profiles of our two cells *i* = 1 and 2. Ideally, the distance would be zero as there should be no difference in the expected location of these cells after normalization. However, the actual distance will be approximately equal to the square root of the sum of 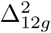 across all (non-DE) genes used in the calculation. This means that the distance between cells with different *s*_*i*_ will be systematically larger than between cells with the same *s*_*i*_. As a result, spurious clusters or trajectories can form (Figure 3) solely due to the log-transformation. Such artificial structures can be highly misleading when characterizing the heterogeneity of a cellular population.

**Figure 3.**
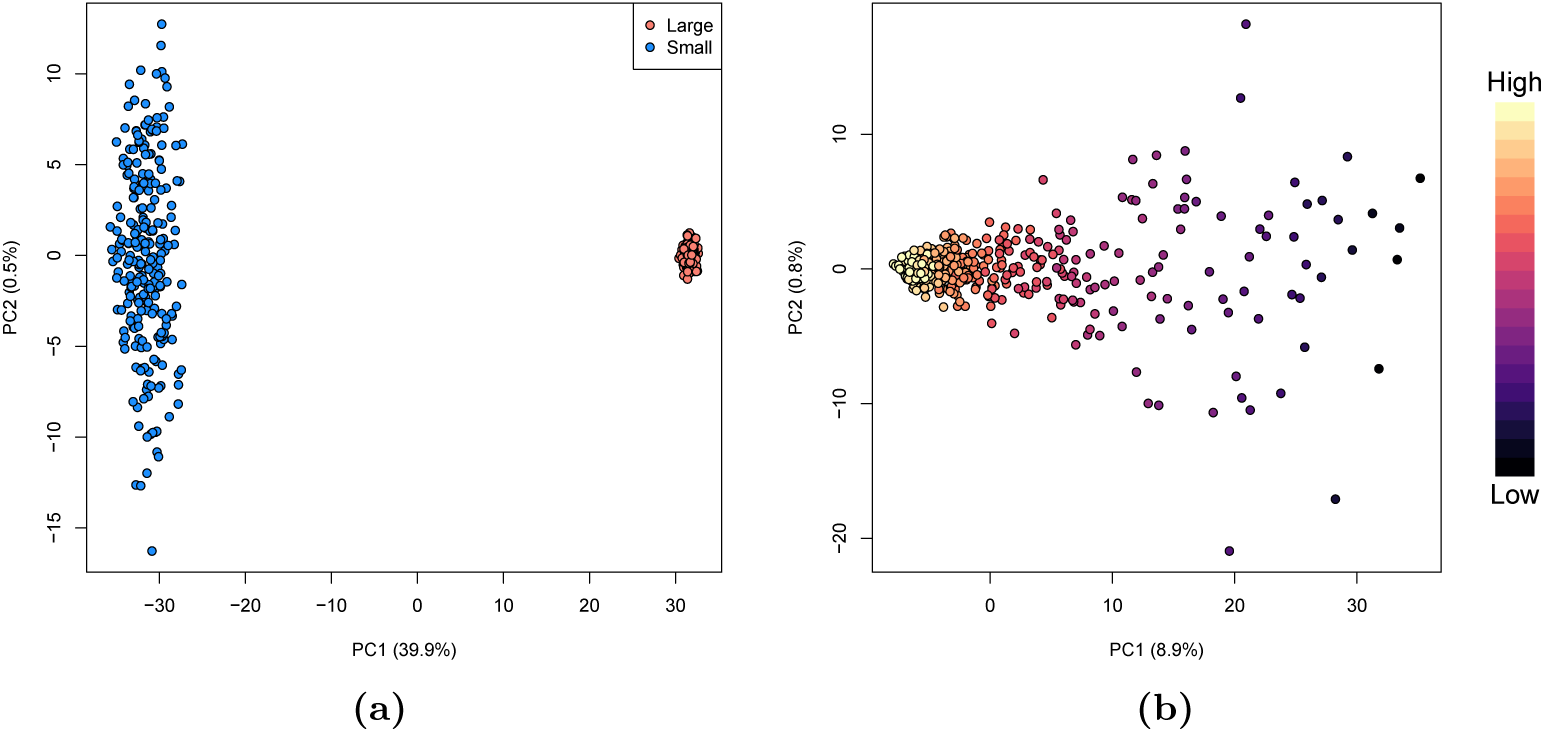
Artificial structures induced by log-transformation in simulations with no true structure. (a) Principal components analysis (PCA) plot of log-normalized expression values for non-DE genes in simulated cells with either very small or very large size factors (see Methods). Each point represents a cell coloured according to its value of *s*_*i*_. The percentage of variance explained by each principal component (PC) is also shown in parentheses. (b) PCA plot for simulated cells with *s*_*i*_ sampled from a continuous distribution. Each point represents a cell coloured by its value of *s*_*i*_.

The discrepancy described in Equation 1 is also consistent with the observations from Hicks *et al.* [1]. *In their study, Hicks et al.* show that the log-normalized expression of many genes is anticorrelated with the “detection rate” (i.e., the percentage of non-zero counts in each cell) in a variety of scRNA-seq data sets. We note that the detection rate is often strongly associated with the size factor due to the sparsity of the data. Any increases to the coverage of a cell (and thus the size factor *s*_*i*_) will naturally increase the detection rate as fewer zeroes are sampled, such that both are correlated with the error from the log-transformation. However, some caution is required when interpreting these correlations in real data due to the conflation of technical effects with genuine biological factors such as, e.g., differences in RNA content across cell types [10, 11].

We wondered whether this problem of artificial differences could be mitigated by using alternative transformations. We considered the square root, which provides variance stabilization for Poisson-distributed counts; and the variance stabilizing transformation (VST) for negative binomial-distributed counts from *DESeq2* [12]. In simulations involving two groups of cells differing only in their size factors, we observed large differences in the group-specific means for non-DE genes after applying each transformation (Figure 4). This mirrors the mean-dependent nature of Δ_12*g*_ and indicates that variance stabilization does not inherently protect against spurious differences. In fact, no transformation can truly stabilize the variance at low counts, as the variance is zero at a mean of zero and must always increase. This is problematic as many scRNA-seq data sets are dominated by low counts – 81% and 89% of non-zero counts are below 5 in the brain and PBMC data sets, respectively – which suggests that favouring a transformation on the basis of its variance stabilization is largely misguided.

**Figure 4.**
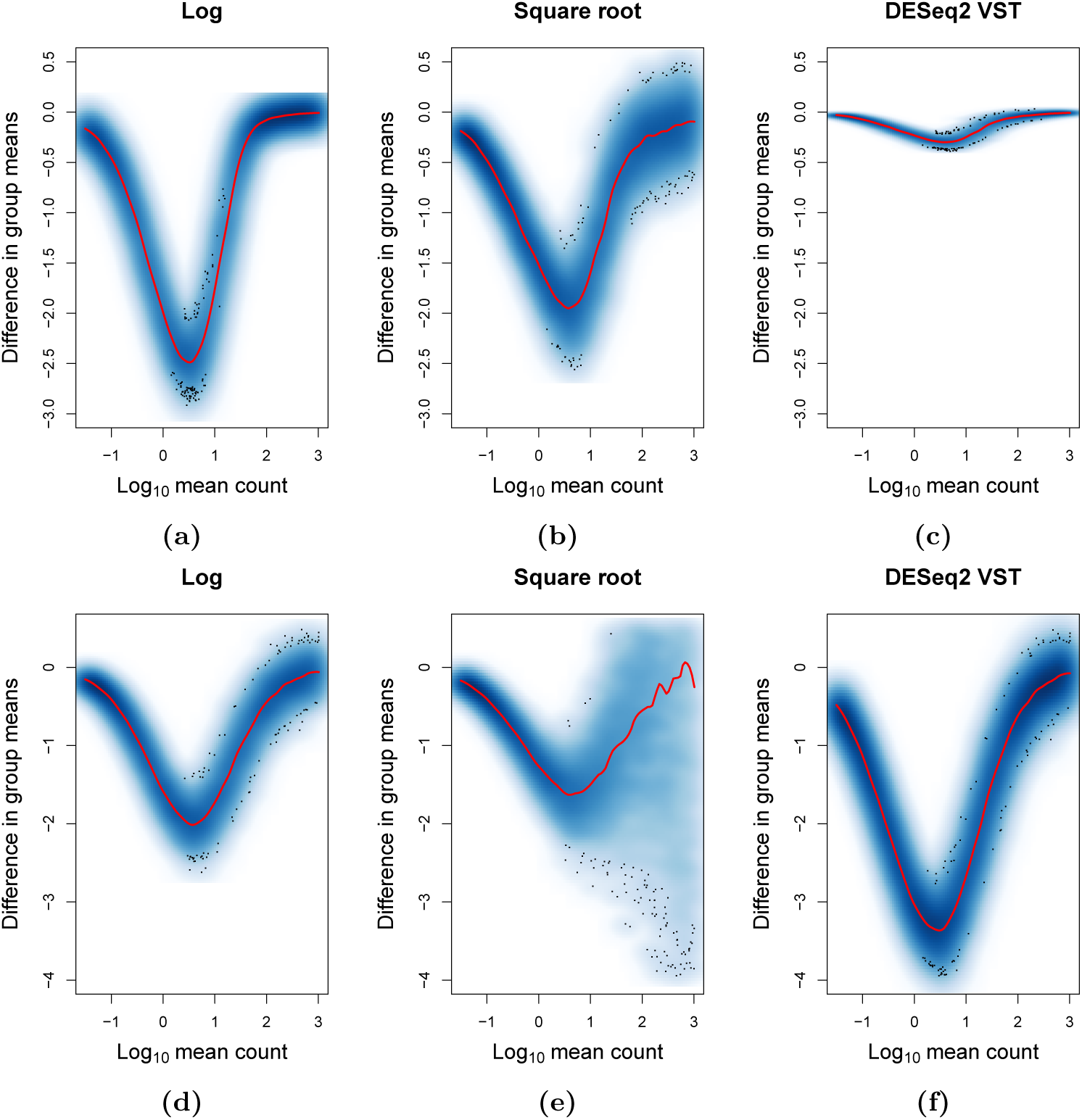
Difference in the group-specific mean transformed expression values against the mean count for each gene, for Poisson-distributed counts (a, b, c) or NB-distributed counts (d, e, f) after applying a variety of transformations (see Methods). Simulated count data was generated for non-DE genes in each of two groups of cells differing only in their size factors (smaller or larger). Colour intensity is proportional to the density of genes, with outliers shown as points. The red line represents a fitted loess curve.

## 3 Validating structure with DE analyses

In our opinion, the effect of greatest concern is the formation of spurious clusters and trajectories. This can easily result in incorrect biological conclusions from an exploratory analysis of a scRNA-seq data set. Fortunately, careful further analysis can provide some protection against these artificial structures. Say that we identify spurious clusters like those in Figure 3. We attempt to characterize these clusters by identifying genes that are differentially expressed between them. Here, the key is to use count-based models like *edgeR* [7] to perform the DE analysis. This directly accounts for complex mean-variance relationships in the count data, thereby avoiding the need for any transformation and concomitant artefacts [13]. If the analysis fails to yield any DE genes, we can infer that the separation was driven by the log-transformation and should be ignored. This strategy is motivated by the presence of differences in the observed effect sizes computed from transformed or untransformed data (Figure 2), possibly leading to different biological conclusions. Our results indicate that untransformed data is more appropriate for these comparisons between groups.

While count-based models provide some protection from spurious structure, it is often the case that we still need to use per-gene transformed values for other steps in the analysis. This includes visualization of expression between groups (e.g., in boxplots, or by colouring dimensionality reduction plots), for which the log-transformation is often used to preserve relative differences across a wide range of expression values. It may also be more convenient to perform DE analyses on the log-transformed values, e.g., using *t*-tests [14, 15], which are faster and simpler to implement than count-based models. Thus, we would like to reduce Δ_12*g*_ as much as possible for these applications. The most obvious method of doing so is to increase the pseudo-count. Indeed, we could force Δ_12*g*_ to zero for all genes by setting *c* to some arbitrarily large value. However, as *c* → *∞*, the log-transformation approaches a linear transformation, i.e.,

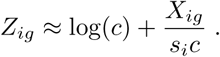

Thus, using an arbitrarily large pseudo-count defeats the intended purpose of the log-transformation. No variance stabilization is achieved and differences in the transformed values cannot be used as proxies for log-fold changes. Instead, we aim to choose a pseudo-count that restricts Δ_12*g*_ to an “acceptable” level for each gene.

## 4 Choosing a larger pseudo-count

We assume that *X*_*ig*_ follows a negative binomial (NB) distribution with mean *s*_*i*_*µ*_*g*_ and dispersion *φ*. This means that we can simplify the expression for Δ_12*g*_ to

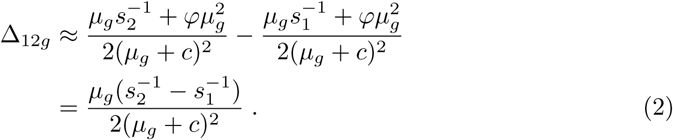

This reaches its maximum absolute value for positive *µ*_*g*_ when *µ*_*g*_ = *c >* 0, yielding

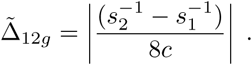

We can control the maximum error 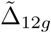 below a threshold *τ* by setting

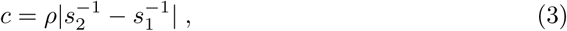

where *τ* = (*ρ*8)^−1^. If we set *ρ* = 1, the maximum difference in the mean log-values should be 0.125 (or [8 log(2)]^−1^ ≈ 0.18, on the log_2_-scale). The observed maximum difference in simulations lies close to this bound at low dispersions (Figure 5). At higher dispersions, the bound becomes less accurate but is nonetheless still conservative.

**Figure 5.**
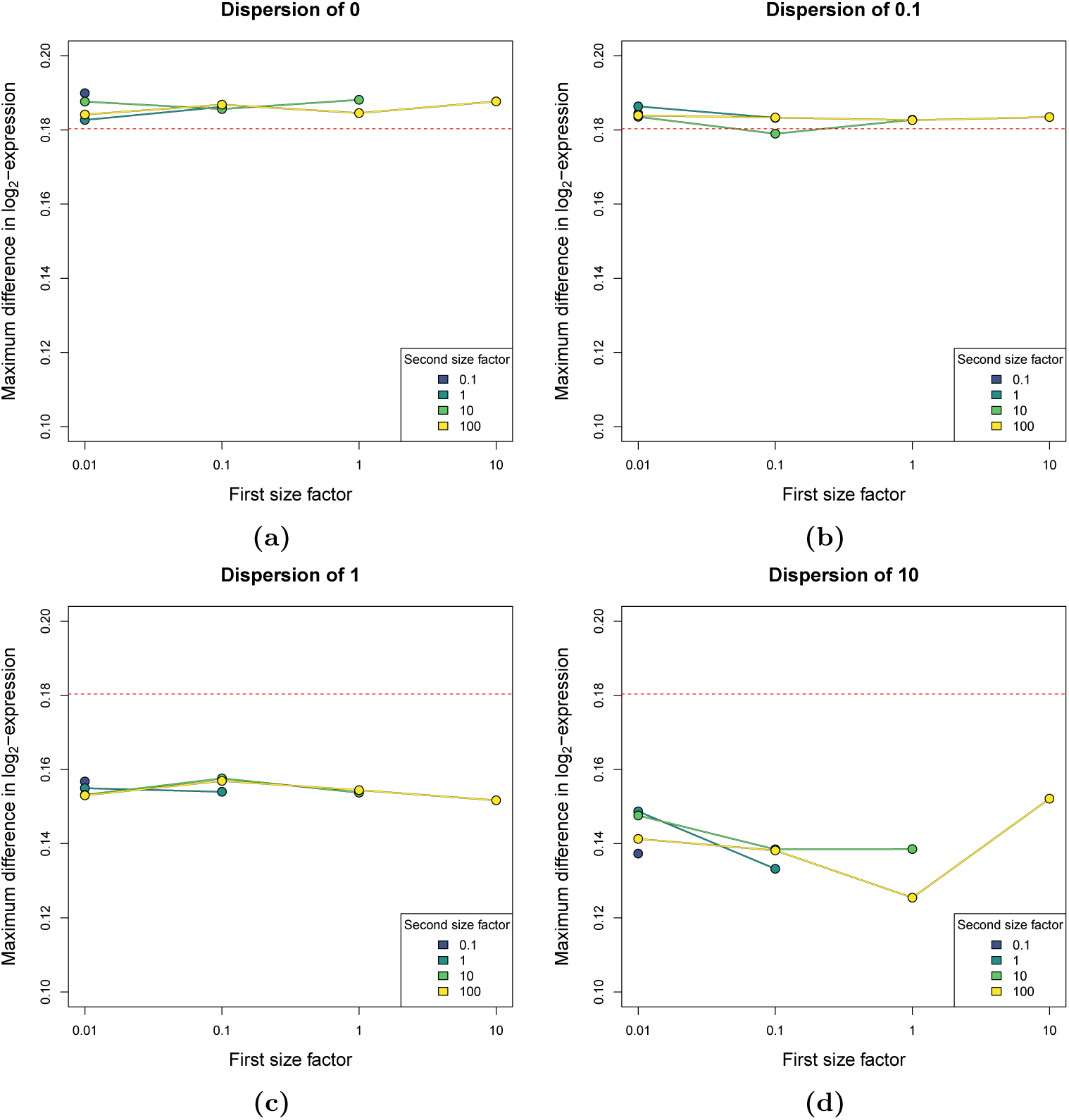
Maximum difference in the log-expression values for non-DE genes between groups of cells differing only in their size factors, after adding the pseudo-count from Equation 3 with *ρ* = 1 prior to log-transformation. Simulations and calculations were performed as described for Figure 1. Results are only shown for scenarios where *s*_1_ *> s*_2_. The red dashed line represents the theoretical bound on the maximum difference.

Equation 3 is most obviously applied to specific comparisons between two groups of cells. In such cases, we can set *s*_1_ and *s*_2_ to the average size factor of each group to control 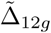 (Figure 6a). However, this may not be generally possible if more than two groups are involved or when groups are not well-defined, e.g., trajectories. The exact choices of *s*_1_ and *s*_2_ are not obvious when calculating a single pseudo-count for the entire data set. We suggest setting *s*_1_ to the smallest size factor and *s*_2_ to the largest size factor (or robust equivalents thereof, e.g., the 5^th^ and 95^th^ percentiles). This ensures that the difference in log-values for a non-DE gene does not exceed the theoretical upper bound for most pairs of cells. We thus define our empirical pseudo-count as

**Figure 6.**
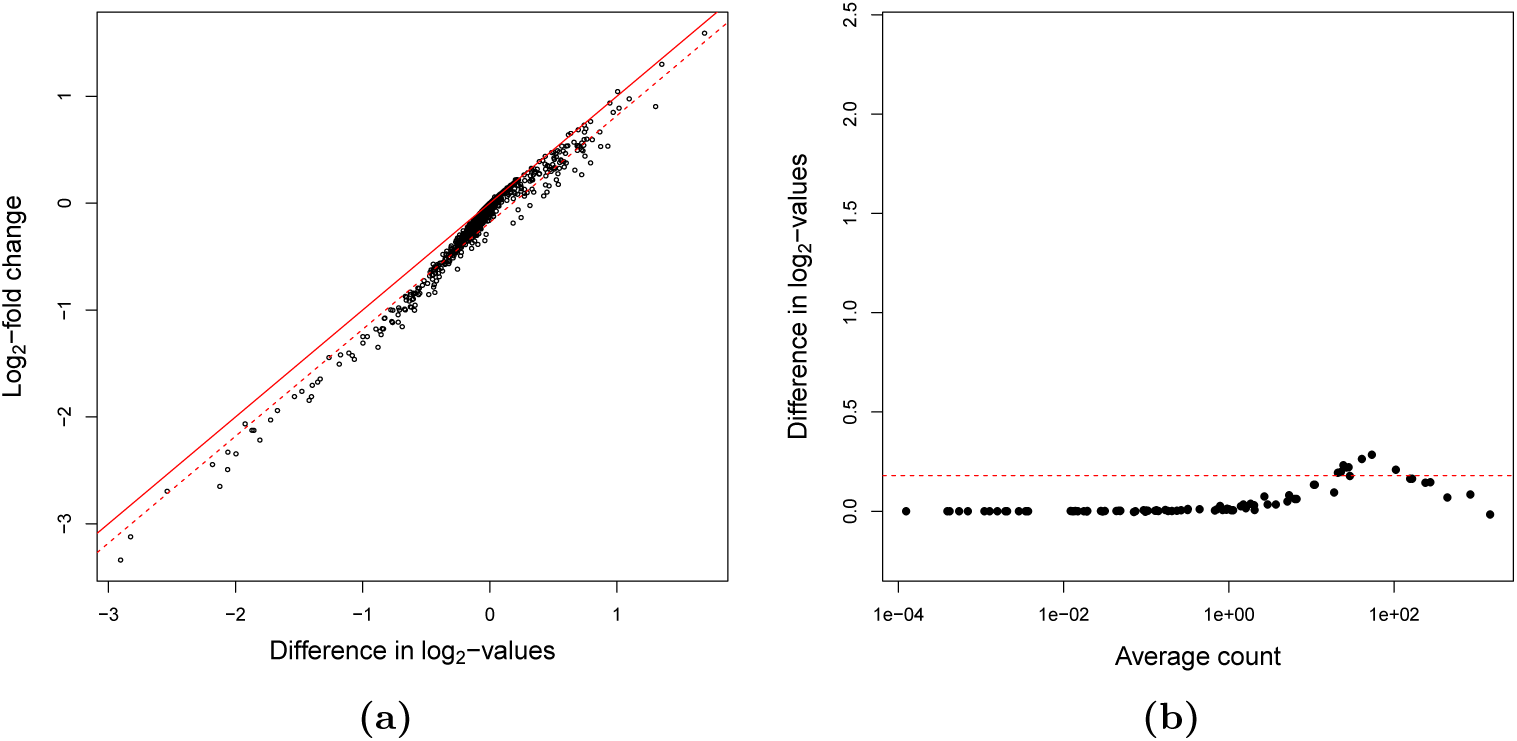
Effect of log-transformation with an increased pseudo-count in real scRNA-seq data. (a) Differences in the mean *Z*_*ig*_ compared to the log-fold change in the PBMC data set, for the same pair of clusters shown in Figure 2d. The pseudo-count was computed using Equation 3 with *ρ* = 1 and using the median size factor for each cluster as *s*_1_ and *s*_2_. The unbroken line represents identity while the dashed line denotes the theoretical limit on 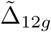. (b) Differences in the mean *Z*_*ig*_ between the top and bottom 20% of libraries with the largest and smallest size factors in the ERCC data set, after adding *ĉ* with *ρ* = 1. The red line denotes the theoretical limit on 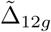.

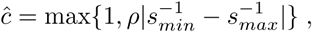

which ensures that the pseudo-count is at least unity when all size factors are equal. We suggest setting *ρ* = 1, which provides a compromise between restricting 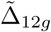 and keeping the pseudo-count as small as possible. Applying this strategy to the ERCC data set limits the maximum error close to the theoretical upper bound (Figure 6b).

We computed *ĉ* for a number of real scRNA-seq data sets to examine its behaviour in practical settings. With *ρ* = 1, we obtained *ĉ* values of 1.36 for the 416B data set [11], 4.05 for the mouse brain data set [9] and 2.22 for the PBMC data set [8]. These values are within the typical range of pseudo-counts used for exploratory analyses of RNA-seq data [16]. They also lie within the range of mean abundances in each data set – we observed 217, 649 and 12636 genes with mean normalized expression values greater than *ĉ* in the PBMC, brain and 416B data sets, respectively. This indicates that the computed *ĉ* is not so large as to entirely negate the variance stabilization from log-transformation. It also justifies our strategy of controlling the maximum spurious difference 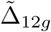, which occurs when the mean normalized expression is close to *ĉ*. (Otherwise, if all genes had lower abundances than *ĉ*, the maximum difference would never be achieved and our procedure would be unnecessarily conservative.)

## 5 Practical use of larger pseudo-counts

We recommend using log-expression values computed with *ĉ* during characterisation of population structures identified from exploratory analyses. This considers the worst-case scenario where there are systematic differences in the size factors between cells in different clusters or along trajectories. (Conversely, it is not required when the distribution of size factors is the same across clusters or trajectories.) By controlling 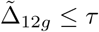, we can mitigate the effects of misleading differences during visualization of expression, as previously discussed. Methods like TREAT [17] can also be applied to test whether the observed log-difference is significantly greater than the theoretical limit *τ.* This provides a statistically rigorous alternative for DE analyses if count-based models are impractical. It is also possible to use log-values computed with *ĉ* in the initial exploratory analysis, which may reduce the risk of detecting spurious structures in the first place. However, this is more difficult to justify as there is no straightforward interpretation for an “acceptable” threshold in the error in the distance between cells.

Pseudo-counts are often set to unity for practical reasons that have little to do with avoiding spurious differences. One reason is to ensure that zero counts always yield zero log-values, which ensures that sparse matrix representations are still effective for reducing memory usage when analyzing large data sets. A related motivation is to ensure that the log-transformed values are always non-negative, which is necessary for procedures such as non-negative matrix factorization. However, there is no need to conflate these considerations with the choice of pseudo-count. Some arithmetic yields

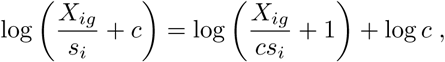

where the constant log(*c*) on the right hand side can be ignored in any step that involves computing differences between log-values. By computing the first term on the right, we can preserve sparsity and non-negativeness in the log-expression matrix.

As an aside, it is also worth commenting on the widespread use of log-transformed counts-per-million (CPM) values and their relatives, e.g., transcripts-per-million or fragments-per-kilobase-million. Log-CPM values are typically computed by adding 1 to the CPM values, for the same reasons mentioned above. However, this is equivalent to using a pseudo-count equal to the library size (i.e., total count across all genes) in millions. For example, data sets with an average library size of 10 million would use a pseudo-count of 10, while data sets with an average library size of 10,000 would use a pseudo-count of 0.01. This is not desirable as data sets with small library sizes are those with the largest Δ_12*g*_ (as *µ*_*g*_ is small) and in most need of large pseudo-counts. In contrast, the approximation of the log-fold change by differences in log-values should improve with greater coverage. Increasing the pseudo-count counteracts this effect such that the log-fold change estimates are not consistent with respect to coverage.

## 6 Discussion

We can speculate on the practical settings in which the log-transformation is most likely to cause distortions. Artifacts are most pronounced when the counts are low and there is large variation in the size factors across cells. Both of these are often observed in data from droplet-based scRNA-seq protocols, as sequencing coverage is shared across a large number of libraries and reaction conditions are not precisely controlled for each droplet. We would also expect the size factors within each experiment to vary across a continuum, yielding artificial trajectories (Figure 3b) rather than distinct clusters (Figure 3a). This suggests that caution is required when interpreting trajectories that are correlated with the size factors, especially in low-coverage droplet-based data.

In the context of scRNA-seq analysis workflows, additional procedures can be used to reduce transformation-induced artifacts. For example, quality control is typically performed to remove cells with small library sizes [18, 19]. This reduces the variation in the size factors and the potential for artificial differences upon log-transformation. In addition, low-abundance genes can be filtered out, which provides further protection as genes with low counts are most affected by the log-transformation. We note that the use of a larger pseudo-count will also reduce the influence of low-abundance genes. This is a more nuanced approach than filtering, as it ensures that strong biological signal in low-abundance genes is not completely discarded for downstream applications.

Data transformations are powerful tools for exploratory analyses of scRNA-seq data. However, they also have the potential to introduce artificial differences, as we have shown for the log-transformation. Our recommendation is to increase the pseudo-count in cases where size factor variation may be responsible for spurious structure in the data.

## 7 Methods

### 7.1 Simulated log-fold changes for non-DE genes

We considered two groups of 10,000 cells where all cells in the same group *j* were assigned the same size factor *s*_*i*_ = *a*_*j*_. For a non-DE gene *g*, the count *X*_*ig*_ for each cell *i* was independently sampled from a negative binomial distribution with mean *s*_*i*_ *µ*_*g*_ and dispersion *φ*. We computed *Z*_*ig*_ for all cells in each group using a pseudo-count *c* of 1 as previously described. The difference in the mean *Z*_*ig*_ between groups represents an estimate of Δ_12*g*_. The maximum difference in the means was then determined by testing a variety of gene abundances *µ*_*g*_ ∈ [10^−3^, 10^3^]. We repeated this procedure across 10 simulation iterations and reported the average of the maximum differences across iterations. This was performed for each simulation scenario defined by the combination of the size factor for two groups (*a*_1_ and *a*_2_, where *a*_1_ *≥ a*_2_) and *φ*.

### 7.2 Log-fold changes in real data sets

We obtained the ERCC data set from the 10X Genomics website (https://support.10xgenomics.com/single-cell-gene-expression/datasets/1.1.0/ercc). We retained all libraries with total counts greater than 100. Size factors were defined as the library sizes, scaled to a mean of unity across libraries. We defined two groups in this data set, with the first group containing the top 20% of libraries with the largest size factors and the second group containing the bottom 20%. For each ERCC transcript, we computed the log-normalized expression values using a pseudo-count of 1. We then computed the differences in the mean log-values between groups to obtain an estimate of Δ_12*g*_. For comparison, we also computed the log-fold change (i.e., *δ*_12*g*_) between groups based on the mean normalized expression. Note that log-fold changes were computed after adding a pseudo-count of 1 to the means of both groups.

For the 416B, brain and PBMC data sets, counts were processed according to a published workflow [19]. Briefly, low-quality cells were removed with the *scater* package [20] and size factors were computed using the deconvolution method [3]. PCA was performed on the log-normalized expression values (computed with a pseudo-count of 1) after modelling the mean-variance trend, and cells were clustered based on their PC scores using the shared nearest neighbours method [21]. In each data set, we identified the two clusters with the largest and smallest median size factors. We computed the difference in the mean log-expression values between clusters and compared this to the log-fold change between clusters calculated from the mean normalized expression. Note that log-fold changes were calculated after adding a pseudo-count of 1 to the means of both groups, such that any discrepancy between the difference in mean log-values and the log-fold change is an estimate of Δ_12*g*_.

### 7.3 Simulations with artifical structures

Counts for 1000 genes in 500 cells were sampled independently from a Poisson distribution with a mean of *λ*_*i*_ for each cell *i*. We set *λ*_*i*_ = 0.1 for half of the cells and *λ*_*i*_ = 10 for the other half. The size factor for each cell was defined as *λ*_*i*_ after being scaled to a mean of unity across cells. We computed log-normalized expression values for all cells with a pseudo-count of 1, as previously described. We then performed PCA and examined the distribution of cells along the first two PCs. We also repeated this simulation where *λ*_*i*_ was sampled from a Uniform(0.1, 5) distribution instead. Note that all genes are non-DE in these simulations, i.e., *E*(*X*_*ig*_*/s*_*i*_) is constant for all *g* and *i*.

### 7.4 Simulations with different transformations

Counts for 10,000 genes in 500 cells were sampled from a Poisson or negative binomial distribution with the mean of *λ*_*i*_*µ*_*g*_ for each cell *i*. We sampled log_2_(*µ*_*g*_) from a Uniform(-5, 10) distribution to obtain a range of gene abundances. We partitioned the cells into two groups of equal size, setting *λ*_*i*_ = 0.1 for all cells in one group and *λ*_*i*_ = 10 for all cells in the other group. For the negative binomial distribution, we used a constant dispersion of 1 for simplicity. The size factor for cell *i* was defined as *λ*_*i*_ after being scaled to a mean of unity across cells. We applied the tested transformations – square root, log, and VST – to the normalized expression values, using a pseudo-count of 1 for the log_2_-transformation and `fitType=“mean”` for the *DESeq2* VST. We then computed the mean of the transformed values in each group for each gene.

### 7.5 Implementation details

All analyses were performed using R 3.5.0 with packages from the open-source Bioconductor project [22]. We used *scater* 1.9.14, *DropletUtils* v1.1.8, *BiocFileCache* v1.5.5 and *DESeq2* v1.21.16. All code is publicly available on GitHub at https://github.com/LTLA/PseudoCount2018.

## 8 Acknowledgements

We would like to thank Dr. John Marioni for helpful suggestions on the manuscript. This work was supported by core funding from Cancer Research UK (award no. A17197 to Dr. Marioni).

